# *Listeria monocytogenes* prophage induction is activated by ppGpp and inhibited by c-di-AMP

**DOI:** 10.64898/2026.05.13.725036

**Authors:** Paarangi Chawla, Yuxing Chen, TuAnh N. Huynh

## Abstract

Bacteriophages, particularly temperate prophages that integrate directly into the host genome, are crucial drivers of bacterial evolution and act as fundamental architects of microbial communities. *Listeria monocytogenes* 10403S has two phage elements – prophage Φ10403S and monocin element. In this study, we found that c-di-AMP, a crucial second messenger in *L. monocytogenes*, regulates phage production. C-di-AMP accumulation down-regulates the gene expression of prophage and monocin gene loci and inhibits phage production, both spontaneous phage production as well as under phage induction through mitomycin-C treatment. We found that in genetically heterogenous cultures, super-infection of non-lysogenic strains with phage-containing strains can significantly amplify spontaneous prophage production. Using these cultures as an induction system, we found other inducers of spontaneous phage production. We found that ppGpp accumulation and nutrient starvation acts as an inducer of the spontaneous prophage production in *L. monocytogenes*. H_2_O_2_ can also play a role in inducing spontaneous phage production. Moreover, Φ10403S prophage production is suppressed in co-cultures of *L. monocytogenes* 10403S with *L. monocytogenes* F2365 and *L. innocua* CLIP11262.

**Importance:** Most *Listeria* and *L. monocytogenes* strains are lysogens, although the impact of phage genes on bacterial host metabolism or host fitness during bacterial competition is still unexplored. Bacteriophages have been shown to influence the evolution of their host and, in several cases, have a major effect on environmental fitness, pathogenicity and/or virulence of bacterial pathogens, either by regulating expression of critical genes or encoding beneficial genes. Prophages provide their hosts with a competitive edge through lysogenic conversion— introducing novel toxins and defense systems—while simultaneously maintaining a molecular trigger, in the form of lysogenic-to-lytic switch, capable of initiating population-wide lysis in response to environmental stress. In the human pathogen *Listeria monocytogenes*, these prophages are near-ubiquitous and studies by other groups have described them as critical regulators of the bacterium’s SOS pathways and its pathogenic lifestyle. This study identifies important novel regulators of phage induction in *Listeria monocytogenes* – c-di-AMP, ppGpp and H_2_O_2_, and how phage induction and production can shape the dynamics of bacterial competition of its host.

## INTRODUCTION

*Listeria monocytogenes* is a Gram-positive foodborne pathogen that is adept at both environmental persistence and adaptation in the mammalian hosts. In its natural niches, *L. monocytogenes* often exists in genetically diverse populations with other strains and *Listeria* species (1, 2). *L. monocytogenes* strains exhibit significant phenotypic heterogeneity in virulence potential, antimicrobial resistance, and stress adaptation, all of which influence infection outcomes (3, 4). Despite this diversity, the mechanisms governing inter-strain and inter-species interactions within these heterogeneous populations remain poorly understood. One potential mechanism for such competition is the induction and releases of infective prophages and tailocins.

Most *L. monocytogenes* strains are lysogens, carrying at least one prophage in their genomes. *L. monocytogenes* strain 10403S, exquisitely well-studied for pathogenesis, harbors two phage-like elements: an F-type tailocin, called monocin; and an infective *Siphoviriade* prophage, called Φ10403S (5). These phage elements are induced by DNA-damaging treatments, such as UV and mitomycin-C, which triggers a *recA*-mediated SOS response (5). Based on well-established model of λ phage activation, upon DNA damage, presumably *L. monocytogenes recA*, binds damaged single-stranded DNA, and cleaves *cI* repressors that are present in both monocin and Φ10403S. Thus, phage production can be induced through DNA-damaging agents like UV irradiation and mitomycin-C. The prophage activation is additionally regulated by other phage-encoded repressors and anti-repressors. Cro- and cI-like repressors are antagonistic transcription factors that form a genetic switch determining whether the phage enters the lytic cycle (*cro*-like repressor) or lysogenic cycle (*cI*-like repressor) (5). *MpaR*, encoded on the monocin element, functions as an anti-repressor by cleaving *cI* repressors of both monocin and Φ10403S (5). *AriS*, encoded in Φ10403S, inhibits SOS response, thereby dampening monocin and prophage induction (6). Finally, transcription of late genes in Φ10403S is also activated by prophage-encoded *llgA*, which is inhibited at 37°C and during *L. monocytogenes* intracellular infection in macrophages, keeping prophage induction low at elevated temperatures and during infection (7).

In addition to UV irradiation and mitomycin-C treatment, *Listeria* prophages can be induced in rich broth cultures by the addition of phosphate, lithium chloride, acriflavine, and nalidixic acid, although the mechanisms are unclear (8). Additionally, during infection of macrophages by *L. monocytogenes*, spontaneous prophage excision occurs in the phagosome through a process called active lysogeny, wherein expression of early phage genes allows prophage excision, but such excision does not lead to the production of infective phages (9). An expanding number of stressors and molecules have been reported to induce prophages, both in SOS-dependent and independent manners, like colibactin, streptozotocin, clinical and environmental xenobiotics like fluoroquinolones (ciprofloxacin), beta-lactams (like ampicillin), ecological stressors like fluctuations in pH, high salinity, metabolic byproducts like short-chain fatty acids (propionic and butyric acids), synthetic chelators like EDTA, etc (5). However, these signals and induction mechanisms haven not been systematically identified for *L. monocytogenes*.

The small nucleotide c-di-AMP is essential for the growth of *L. monocytogenes* in rich broth media and during infection. *L. monocytogenes* synthesizes c-di-AMP by a single diadenylate cyclase, *dacA* (also called *cdaA*), and hydrolyzes it by the phosphodiesterases *pdeA* (also called *gdpP* in other bacteria) and *pgpH* (10, 11). While *dacA* mutants have been studied extensively in different bacteria, the Δ*pdeA* Δ*pgpH* mutant (hereafter denoted as ΔPDE), which accumulates c-di-AMP, has proven to be a particularly useful genetic tool to define the function of c-di-AMP. This mutant grows normally in broth media, but is highly sensitive to stress, like osmotic stress and cell-wall targeting antibiotics, and attenuated for virulence.

To profile the molecular defects conferred by c-di-AMP accumulation, we previously profiled the transcriptomes of the wild-type (WT) and ΔPDE strains during growth in a defined medium, called *Listeria* Synthetic Medium (LSM) (5). Compared to WT, the ΔPDE strain was significantly down-regulated mostly just two clusters of genes – *prfA*-regulated virulence genes, and genes within the Φ10403S prophage and monocin clusters (12). In this study, we found the ΔPDE strain to be defective for the production of infective Φ10403S phages, both under mitomycin-C treatment and spontaneously in stationary phase cultures. Spontaneous prophage induction in the WT strain was driven by both ppGpp and H_2_O_2_, and phage induction defect in the ΔPDE strain was partially due to the lack of ppGpp. Interestingly, Φ10403S prophage production is suppressed in co-cultures *with L. monocytogenes* F2365 *and L. innocua* CLIP11262, suggesting that metabolites produced during inter-bacterial competition can modulate prophage induction, potentially shaping the population dynamics of *Listeria* in the environment.

## RESULTS

### c-di-AMP accumulation inhibits phage production in *L. monocytogenes*

We previously found that the ΔPDE strain, which accumulates c-di-AMP, significantly down-regulates the prophage and monocin gene loci when grown in *Listeria* Synthetic Medium (LSM), compared to the wild-type (WT) strain (**Fig. S1**). Specifically, within these clusters, the expression of regulatory genes was comparable between WT and ΔPDE strain, but the expression of early and late genes, encoding phage structural elements, was down-regulated by at least 10-fold in ΔPDE compared to WT. Since Φ10403S prophage can be induced to make infective phages using mitomycin-C and UV irradiation, we queried the effect of this transcriptional down-regulation on phage production. Under mitomycin-C treatment, the WT and ΔPDE strains exhibited similar lysis kinetics (**Fig. S2**) and produced increasing phage particles at similar rates. However, at any given time point, ΔPDE consistently produced 1-2 log fewer phages than WT, with the most pronounced defect at up to 6 hours post treatment (**Fig. 1A**). Combined, these data suggest that the ΔPDE strain may exhibit aberrant activities of phage regulators and a defect in the production of phage particles. Since phage production is decreased in ΔPDE strain at several different time points, this defect phenotype does not seem to be due to premature lysis of ΔPDE strain.

**Figure 1:**
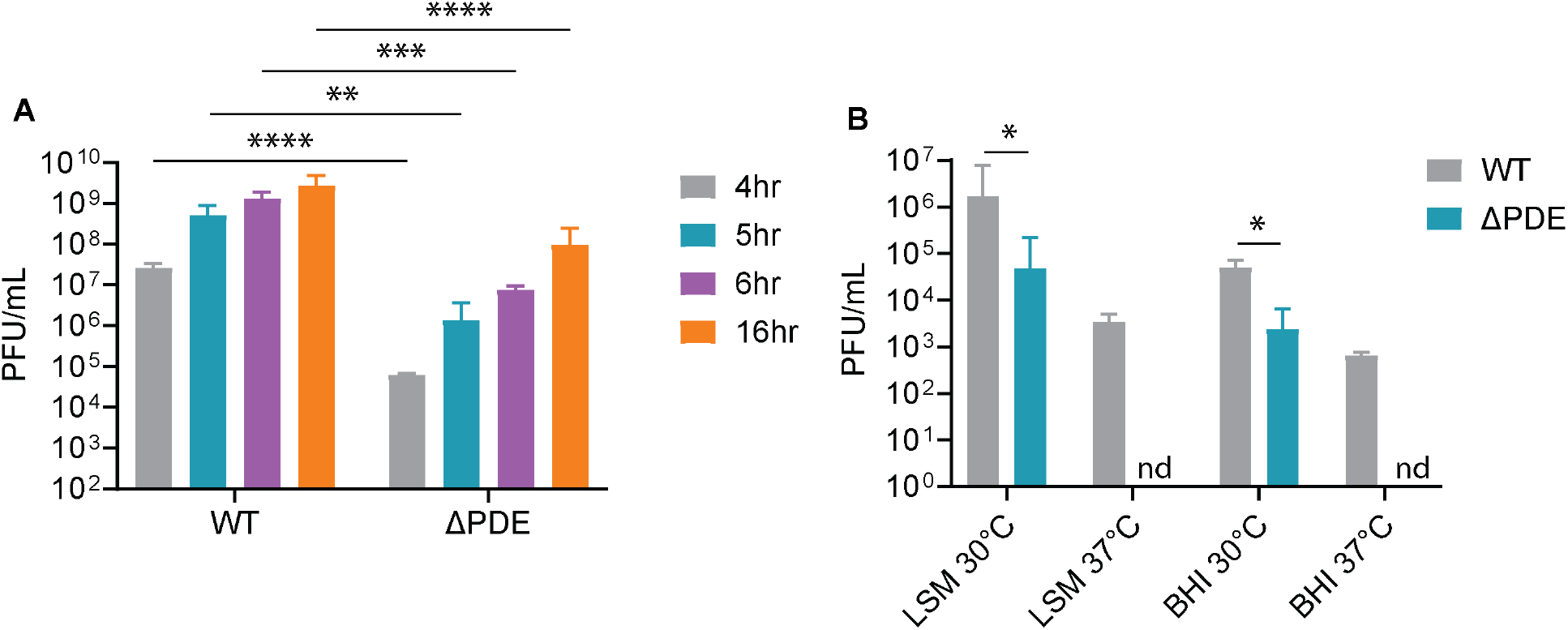
*L. monocytogenes* 10403S spontaneously produces infective phages and the ΔPDE strain is defective for phage production. **A**. Phage production under mitomycin-C treatment. Cultures were grown in LSM to the same density, treated with mitomycin-C for indicated time, and culture supernatants were used to infect *L. monocytogenes* MACK861 strain to quantify phage particles. **B**. Cultures were grown in at 30°C in LSM or BHI statically, 37°C in LSM or BHI with shaking for 24 hours. Infective phages were quantified using *L. monocytogenes* MACK861 strain infection and plaque quantification. Detection limit was 1 PFU/mL. Data are average of at least three independent experiments. Error bars represent standard deviations. Statistical analyses were performed by Student’s t-test between indicated pairs. *, P < 0.05; **, P < 0.01; ***, P < 0.001; ****, P < 0.0001; nd, not detected.

Osmotic stress has been shown to induce the excision of lambdoid prophages in *E. coli* and *Staphylococcus aureus* by activating the SOS response, just like mitomycin-C (13, 14). The primary response to osmotic stress is potassium (K^+^) uptake, which is inhibited by c-di-AMP (15). If the prophage induction defect of ΔPDE is due to potassium uptake defect, then it should be exacerbated with limiting potassium levels and rescued with excess potassium. However, we found potassium availability to have negligible effects on prophage production by WT and ΔPDE or the difference between these strains (**Fig. S3**).

Given that our RNAseq analysis was performed for cultures under normal growth in LSM, we examined if the down-regulation of prophage genes is physiologically relevant by determining spontaneous Φ10403S prophage induction. We found that *L. monocytogenes* can produce significant levels of infective phages in broth cultures spontaneously, without a DNA-damaging treatment. In a defined *Listeria* Synthetic (LSM) medium at 30°C, WT produced up to ~ 5×10^6^ pfu/mL of infective phages (**Fig. 1B**), compared to ~ 5 × 10^9^ pfu/mL as a result of mitomycin C treatment (**Fig. 1A**). Phage production was detectable at early time points, but accelerated at late growth phase (**Fig. S4**). In addition, WT produced ~10^5^ pfu/mL under normal growth in the rich medium Brain Heart Infusion (BHI) broth at 30°C (**Fig 1B**). Although the expression of late phage is largely inhibited at 37°C (7), WT still produced ~ 5×10^3^ pfu/mL in LSM and ~ 6×10^2^ pfu/mL in BHI at 37°C (**Fig 1B**).

Compared to WT, the ΔPDE was defective for phage production in all conditions, despite minimal growth defect (**Figs. 1A, 1B** and **S5**). Most notably, we did not detect phages produced by ΔPDE at 37°C, suggesting that prophage repressors are more active in this strain.

### Super-infection and some intra-species competition amplify spontaneous prophage production

Having found consistent Φ10403S phage particle production by *L. monocytogenes*, we next queried the impacts of phage release on *L. monocytogenes* population. We found that Φ10403S phages did not affect the growth of a WT lysogen strain, even at a high multiplicity of infection (MOI) of 1 but readily lysed the ΔΦWT non-lysogenic strain following infection even at a low MOI of 0.01 (**Fig. S6**).

Following infection with Φ10403S phage particles, both WT and ΔPDE lysogen strains continued to produce infective phages. Despite poor growth and rapid lysis following this treatment, phage-cured strains readily produced infective Φ10403S, resulting in ~3.5-log higher titers than their respective lysogen strains (**Fig. 2**). Interestingly, whereas ΔPDE produced fewer phages than WT, the ΔΦΔPDE strain was indistinguishable from ΔΦWT, indicating that a prophage-encoded regulator inhibits phage induction and production in ΔPDE.

**Figure 2:**
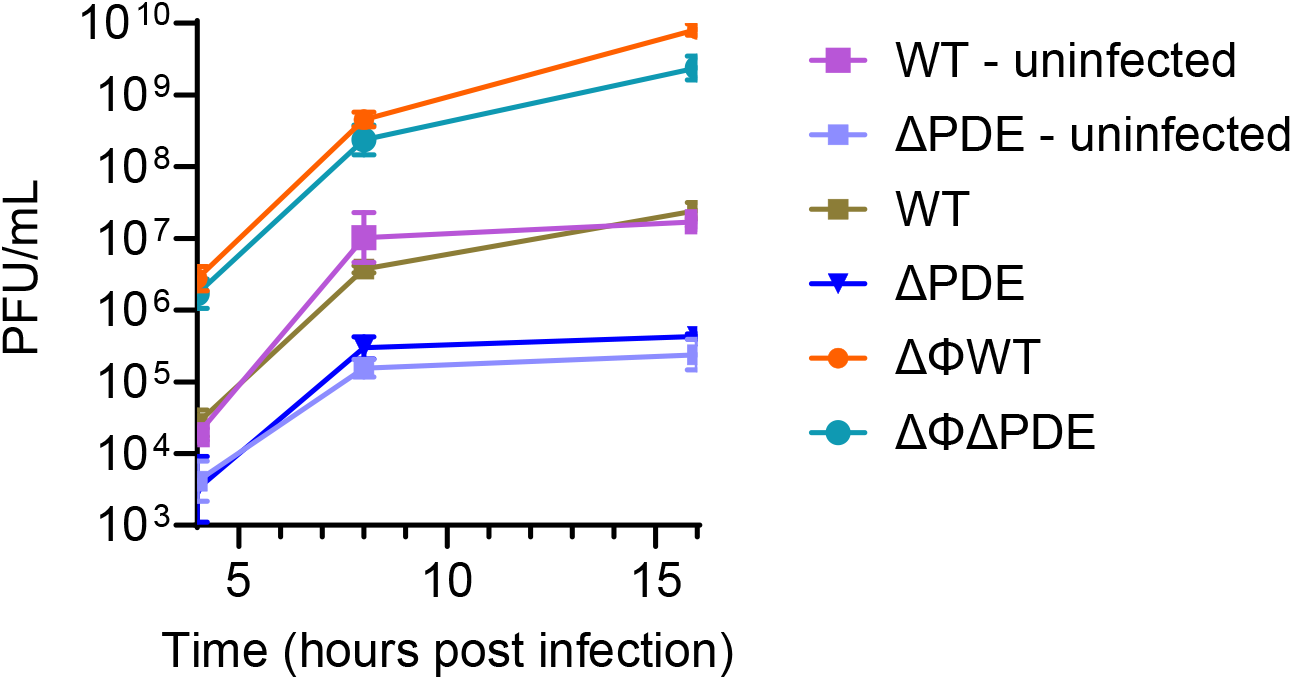
Φ10403S virion production and release are amplified after phage infection in *L. monocytogenes* 10403S. *L. monocytogenes* was grown in LSM to mid-log, then remained uninfected or was infected with Φ10403S phage particles at MOI 0.1. The resulting phage production was quantified from culture supernatants at indicated time point post infection. Data are average of at least three independent experiments. Error bars represent standard deviations.

The increased sensitivity of phage-cured *L. monocytogenes* to infective phage particles was expected, because prophages are known to confer immunity to phage infection. Nevertheless, this observation could be exploited to experimentally generate robust spontaneous phage production, to facilitate mechanistic investigation of prophage induction. To this end, we quantified phage production by co-cultures of lysogen and phage-cured strains. Indeed, the WT + ΔΦWT and ΔPDE + ΔΦWT co-cultures produced nearly 10^10^ pfu/mL of infective phages, almost higher than phage levels under mitomycin C treatment (**Fig. 3A**). The ΔPDE + ΔΦΔPDE culture produced fewer phages than those pairs, but still at ~2.5-log more phages than ΔPDE alone.

**Figure 3:**
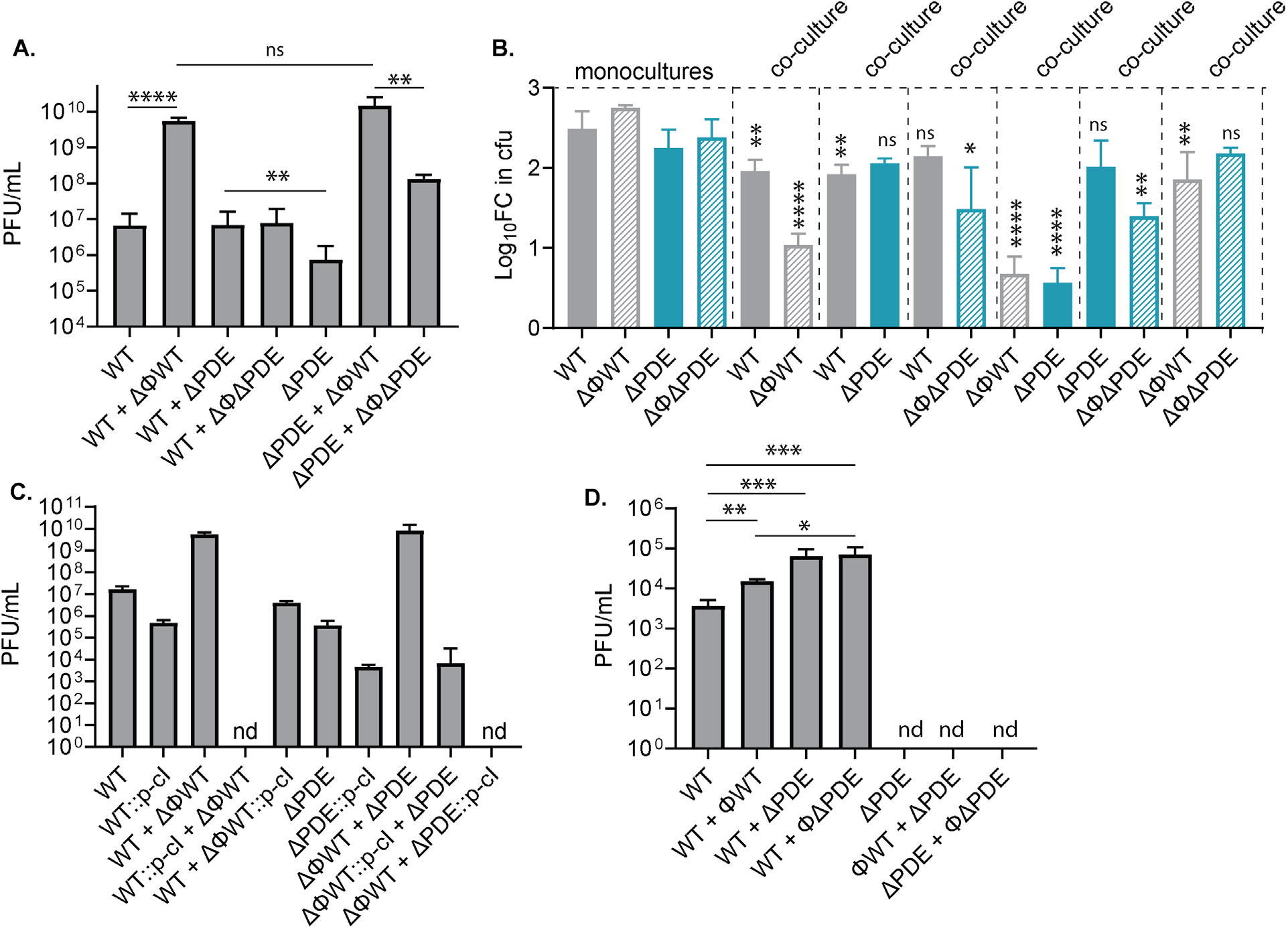
Spontaneous prophage induction is significantly amplified in co-cultures, compromising bacterial growth. **A**. Phage production by *L. monocytogenes* cultures grown statically in LSM at 30°C for 24 hours, quantified as plaque formation following infection of *L. monocytogenes* MACK861 strain. **B**. Bacterial growth in LSM at 30°C, quantified from experiments performed in (A) as log_10_fold change in cfu between 0 hour and 24 hour. **C**. Phage production by *L. monocytogenes* cultures of *cI*-over-expressing strains grown statically in LSM at 30°C for 24 hours, quantified as plaque formation following infection of *L. monocytogenes* MACK861 strain. **D**. Phage production by cultures grown in LSM at 37°C with shaking for 24 hours; nd: not detected; ns, not significant. Detection limit was 1 PFU/mL. Data are average of at least three independent experiments. Error bars represent standard deviations. Statistical analyses were performed by one-way ANOVA with Dunnett’s multiple comparisons. Statistical comparisons in C are between indicated strains and corresponding monocultures. *, P < 0.05; **, P < 0.01; ***, P < 0.001; ****, P < 0.0001; nd, not detected; ns, not significant.

In these co-cultures, high phage production generally compromised bacterial growth, but especially of the phage-cured strains, as anticipated (**Fig. 3B**). Consistent with its remarkably high phage titer, the ΔPDE + ΔΦWT co-culture appeared to clear at the point of phage harvest, indicating bacterial lysis. As a result of lysis, ΔΦWT and ΔPDE strains exhibited limited growth in a co-culture (**Fig. 3B**). Interestingly, although the WT + ΔΦWT co-culture produced a similar phage titer, this pair maintained a relatively stable culture, with significant growth by both strains. Because the ΔPDE culture did not lyse following super-infection with Φ10403S phage particles, its lysis in a co-culture with ΔΦWT suggests that there are additional bactericidal factors released by ΔΦWT.

Phage repressors play a primary role in phage resistance. The Φ10403S prophage encodes a *cI*-like repressor that inhibits phage induction and subsequent phage production. In the WT + ΔΦWT and ΔPDE + ΔΦWT co-cultures - the highest phage-producers, ectopic expression of *cI*-like repressor in ΔΦWT strain significantly reduced phage titers to match WT and ΔPDE monocultures, confirming that superinfection of phage-cured strains amplifies phage production in a genetically heterogeneous population (**Fig. 3C**). Strikingly, over-expression of *cI*-like repressor in the lysogen strains completely abolished phage production by these co-cultures, although it still allowed phage production by the WT and ΔPDE monocultures (**Fig. 3C**).

Finally, a boost in prophage induction occurred not only in phage-cured and lysogen strain pairings, but also in WT + ΔPDE co-culture during growth at 37°C. Interestingly, while neither ΔPDE nor ΔΦΔPDE produced phages under this condition, both strains appeared to increase phage production by WT in co-cultures (**Fig. 3D**).

### ppGpp is a driver of spontaneous prophage induction in *L. monocytogenes*

While it is unsurprising that co-cultures of lysogen and phage-cured strains produce high levels of phage particles, they provided a robust induction system to investigate what signals induce prophages under normal growth. Given that phage titers were higher in the defined medium LSM than the rich medium BHI, and that phage production accelerated in late growth phase, nutrient depletion could contribute to phage induction. To test this, we quantified phage production in LSM with reduced glucose or amino acid contents. These modifications did not affect monocultures, but significantly increased phage production by WT + ΔPDE (**Fig. 4A**). The ΔPDE + ΔΦWT pair was not affected, likely because it already produced very high phage levels which caused significant bacterial lysis in the co-culture.

**Figure 4:**
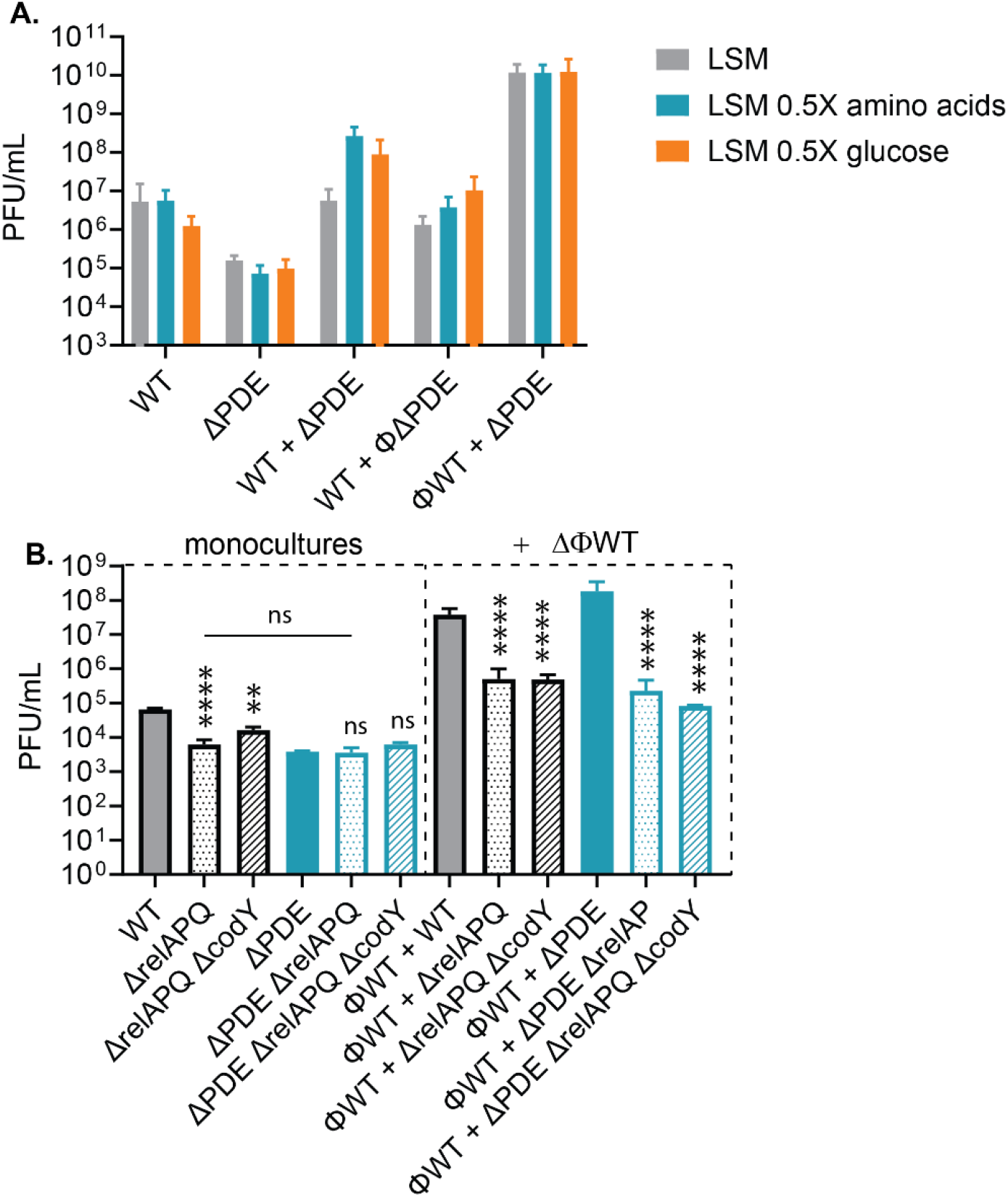
Nutrient depletion and ppGpp accumulation triggers Φ10403S prophage induction. **A**. Phage production by *L. monocytogenes* following 24 hours of static growth at 30°C, in LSM or modified LSM as indicated. **B**. Phage production following 24 hours of static growth in BHI at 30°C. BHI was chosen because Δ*relAPQ* strains do not grow in LSM. Data are average of three independent experiments. Error bars represent standard deviations. Statistical analyses were performed by one-way ANOVA with Dunnett’s multiple comparisons. In panel B, comparisons were made with the first strain in each group. *, P < 0.05; **, P < 0.01; ***, P < 0.001; ****, P < 0.0001; ns, not significant.

Amino acid starvation triggers the accumulation of ppGpp, synthesized by *relA, relP*, and *relQ* in *L. monocytogenes*. Deletion of *relAPQ* not only reduced phage titers in WT, but also diminished the difference between WT and ΔPDE backgrounds. Furthermore, *relAPQ* deletions significantly reduced phage production by these strains in the presence of ΔΦWT (**Fig. 4B**). These data suggest that ppGpp is an initiator of Φ10403S prophage induction, especially in co-cultures, and a lack of ppGpp contributes to ΔPDE phage induction defect.

In low GC Gram-positive bacteria such as *L. monocytogenes*, ppGpp accumulation results in repression of *codY* gene, which transcriptionally represses branched chain amino acid synthesis. This *codY* repression is relieved in Δ*relAPQ* strains. Deletion of *codY* partially rescues some of Δ*relAPQ* phenotypes, such as virulence attenuation. We found that Δ*relAPQ* and Δ*relAPQ codY*::spec alleles resulted in similar phage titers, indicating that *codY* does not mediate prophage induction by ppGpp (**Fig 4B**).

### H_2_O_2_ also contributes to spontaneous prophage induction in *L. monocytogenes*

Reactive oxygen species, particularly H_2_O_2_, is a known inducer of prophage excision (16, 17). Using the fluorescent dye H_2_DFCA as a reporter of reactive oxygen species, we found a remarkably high fluorescence signal in the ΔPDE + ΔΦWT co-culture (**Fig. S7A**). However, H_2_DFCA has recently been shown to be inaccurate in reporting reactive oxygen species in *E. coli* (16, 17). As a complementary approach to evaluate the role of H_2_O_2_ in spontaneous phage induction in *L. monocytogenes*, we queried the effect of catalase (*kat*) over-expression on phage titers. First, *kat* gene over-expression significantly increased H_2_O_2_ resistance, verifying its catalase function (**Fig. S7B**). In WT and ΔPDE monocultures, *kat* over-expression both reduced phage titers by ~1 log, but did not abolish the difference between these two strains. In WT + ΔΦWT and ΔPDE + ΔΦWT co-cultures, the effect of increased catalase activity on phage production was modest (**Fig. 5**). This data indicates a minor role for H_2_O_2_ in spontaneous prophage induction in *L. monocytogenes*.

**Figure 5:**
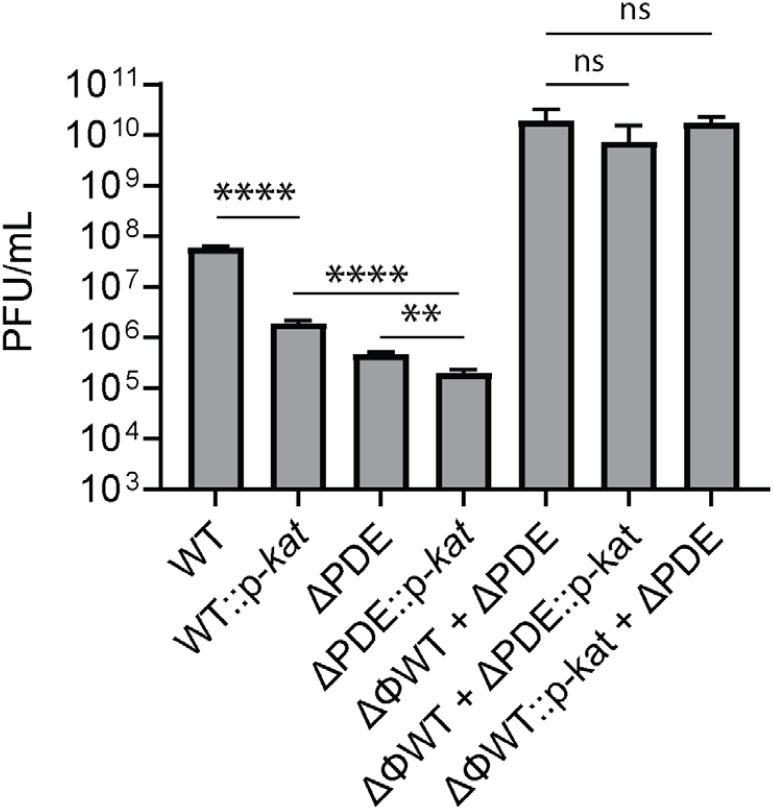
H_2_O_2_ contributes to spontaneous Φ10403S prophage induction. *L. monocytogenes* cultures were grown in LSM at 30°C for 24 hours. Phages were quantified following infection of *L. monocytogenes* MACK861 strain. Data are average of at least three independent experiments. Error bars represent standard deviations. Statistical analyses were performed by Student’s t-test between indicated pairs. *, P < 0.05; **, P < 0.01; ***, P < 0.001; ****, P < 0.0001; ns, not significant.

### Competition with other *Listeria* species suppresses prophage induction by *L. monocytogenes* 10403S

In the environment, *L. monocytogenes* exists in heterogeneous populations with other *L. monocytogenes* strains and *Listeria* species. We therefore examined if Φ10403S prophage induction could occur in such mixed populations, and its impacts on population dynamics. We assembled co-cultures of strain 10403S (serotype 1/2a, clonal complex 9) with *L. monocytogene* strain F2365 (serotype 4b, clonal complex 1), or with *L. innocua* CLIP11262. Interestingly, both *L. monocytogenes* F2365 and *L. innocua* CLIP11262 suppressed Φ10403S prophage induction by *L. monocytogenes* 10403S (**Fig. 6A**). Nevertheless, lysogen and phage-cured 10403S strains grew similarly, and their growth was also not impacted by the presence of other *Listeria* strains (**Fig. 6B**).

**Figure 6:**
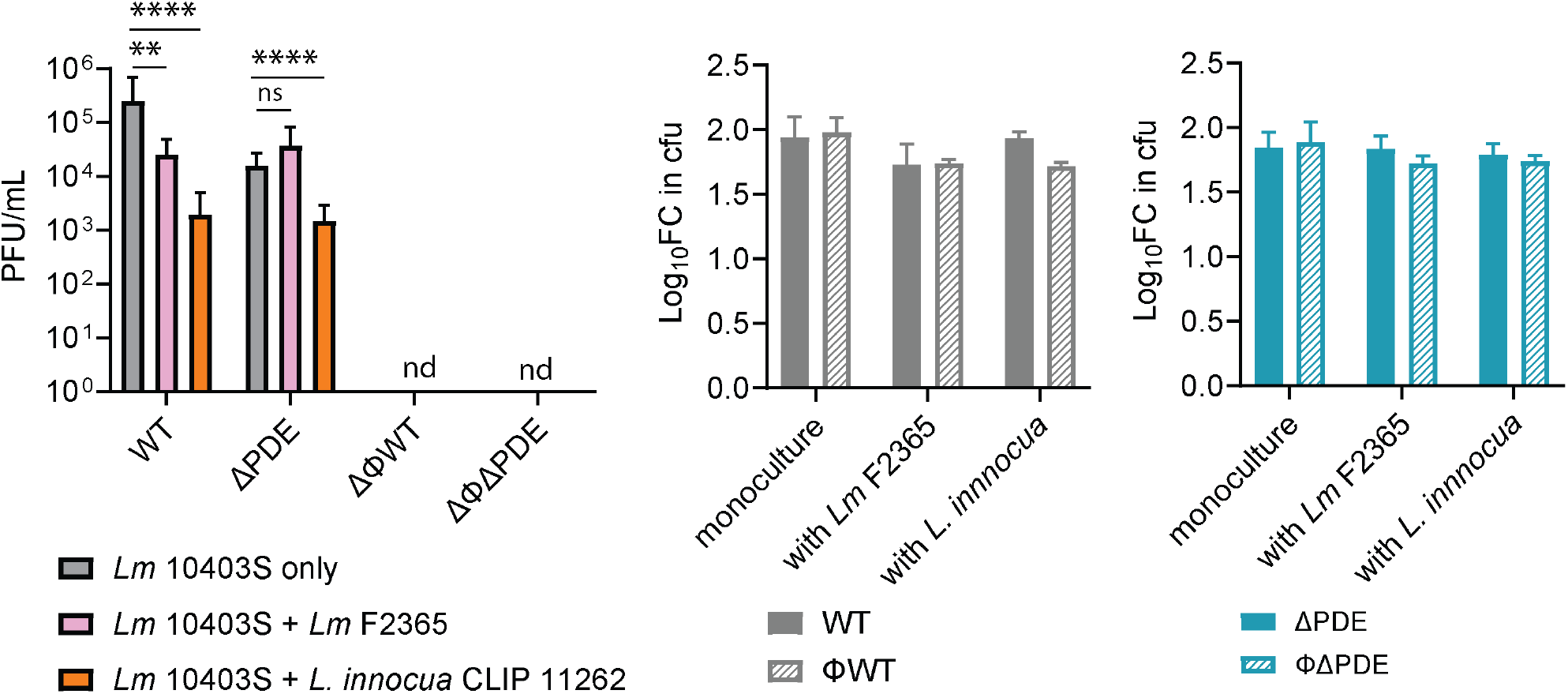
Competition with other *Listeria* strains suppresses Φ10403S prophage induction in *L. monocytogenes* 10403S. Φ10403S prophages, harvested from co-culture supernatants after 24 hours of static growth in BHI at 30°C, were used to infect *L. monocytogenes* MACK861 strain to quantify phages. The ΔΦWT and ΔΦΔPDE strains did not produce detectable phages in monocultures or co-cultures. Detection limit is 1 PFU/mL. **B**. Growth rates of *Listeria monocytogenes* 10403S in monocultures and in co-cultures with *Listeria monocytogenes* F2365 and *Listeria innocua* CLIP 11262. Lysogen and phage-cured 10403S strains grew similarly, and their growth was also not impacted by the presence of other *Listeria* strains. Data are average of three independent experiments. Error bars represent standard deviations. Statistical analyses were performed by one-way ANOVA with Dunnett’s multiple comparisons. *, P < 0.05; **, P < 0.01; ***, P < 0.001; ****, P < 0.0001; nd, not detected.

## DISCUSSION

*Listeria* species are ubiquitously present in different natural environments, often in genetically heterogeneous communities. Only *Listeria monocytogenes* infects humans, but its diverse strains exhibit vastly different virulence potential (3, 18). Furthermore, there appears to be a fitness trade-off between virulence and environmental persistence. For example, hypo-virulent strains are enhanced for biofilm formation and stress resistance, whereas hyper-virulent strains can exhibit growth and stress response defects in broth cultures (19, 20). Defining the mechanisms underlying *Listeria* population dynamics is therefore important to understand the selection for hyper-virulent strains.

Most *Listeria* species and *L. monocytogenes* strains are lysogens and contain more than one prophage or phage elements, although the impact of phage genes on bacterial host metabolism is not yet determined. Nevertheless, prophages are considered an important determinant of *Listeria* population dynamics, as the switch of lysogens to a lytic cycle lyses the host and can release infective phages that attack other bacteria. Therefore, we considered Φ10403S prophage induction in monocultures and mixed co-cultures of *L. monocytogenes*. A previous study reported that *Listeria* prophages can be spontaneously induced to produce phage particles in certain BHI broths (7). Extending those observations, we found that Φ10403S is spontaneously induced under normal growth in various broth culture conditions, both in the defined medium LSM and rich medium BHI, and at temperate temperature of 30°C and elevated temperature of 37°C. Co-cultures of mixed *L. monocytogenes* 10403S strains generally enhance prophage induction significantly.

Additional analyses suggest that nutrient depletion and the stringent alarmone ppGpp is an inducer of spontaneous Φ10403S induction. First, we observed consistently higher phage titers in the defined medium LSM than in the rich medium BHI, and phage production accelerated in late growth phase. Furthermore, in the WT *L. monocytogenes* 10403S strain, we found lower phage production by the ppGpp^0^ (Δ*relAPQ*) strain. Although the effect of ppGpp in monocultures was rather modest, it played a significant role in co-cultures of mixed *L. monocytogenes* 10403S strains, underlying bacterial competition for nutrients and consistent with the effect of nutrient depletion on prophage induction in other ecological niches. In Gram-positive bacteria, ppGpp accumulation is triggered by branched chain amino acid starvation, and as a result depletes GTP pool. A major target of ppGpp accumulation and GTP depletion is *codY*. The GTP-bound form of *codY* represses the expression of genes encoding branched chain amino acid synthesis. Under ppGpp accumulation and GTP depletion, *codY* repression is relieved (7). Consistent with the phage-inducing effect of ppGpp, amino acid depletion significantly increased phage titer in some *L. monocytogenes* co-cultures. Interestingly, while other metabolic and virulence defects of ppGpp^0^ strains are suppressed by *codY* deletion (21–23), the prophage inducing defect of ppGpp was independent of *codY*, raising interesting questions about what molecular targets mediate phage induction and production under stringent response.

Reactive oxygen species, such as H_2_O_2_ and hypochlorite, are known inducers of prophages, although such induction does not necessarily result in production of phage particles (16, 17). Furthermore, bacterial competition has been shown to generate reactive oxygen species (24). Indeed, using the fluorescence dye H_2_DCFDA as a ROS reporter, we found a significant fluorescence signal in the highest phage-producing culture (ΔΦWT + ΔPDE). We acknowledge that the accuracy of H_2_DCFDA as a ROS reporter in *L. monocytogenes* has not been determined. While signal generated by this dye correlates with oxidative stress response gene expression (24), it appears to also respond to non-ROS metabolites in *E. coli* (25). In our experimental set up, ROS is unlikely to be the sole inducer of Φ10403S prophage. We observed phage production in BHI, which is rich in glutathione, a highly effective antioxidant that can be efficiently imported into *L. monocytogenes* (26). Nevertheless, we found a significant effect of catalase over-expression on spontaneous phage induction.

While we observed significant amplification of Φ10403S prophage induction in mixed *L. monocytogenes* 10403S strains, the presence of *L. monocytogenes* F2365 and *L. innocua* CLIP11262 surprisingly suppressed Φ10403S phage production, both in defined and rich media. This suppression cannot be explained by nutrient depletion or oxidative stress, since BHI is a rich media and is rich in glutathione, an antioxidant. Further studies are warranted to investigate prophage induction in bacterial communities. Nevertheless, prophage induction did not appear to compromise *Listeria* host fitness, as lysogen and phage-cured 10403S strains grew similarly in competition with other Listeria.

The second messenger c-di-AMP is a global regulator of bacterial stress response and pathogenesis. Although c-di-AMP is essential for *L. monocytogenes* growth in rich broth media, its accumulation attenuates virulence and confers sensitivity to several stress conditions, such as osmotic and cell wall stresses (7). Here we found that c-di-AMP accumulation in the ΔPDE strain also inhibits Φ10403S prophage induction, both spontaneously under normal growth and mitomycin-C induction. Under normal growth, ΔPDE expressed wild-type levels of phage regulator genes, but significantly down-regulated phage structural genes. Under mitomycin-C treatment, ΔPDE produced consistently fewer phages than WT at any examined time point but exhibited similar rates of phage production to WT. Super-infection of WT and ΔPDE lysogen strains also resulted in significantly fewer Φ10403S phage titers from ΔPDE, but infection of phage-cured strains resulted in similar phage titers. These data suggest that c-di-AMP accumulation impedes the activity of a phage regulator encoded in the Φ10403S prophage, such as *cI, ariS*, or *llgA*. We noticed that phage induction was completely abolished in the ΔPDE strain at 37°C, a condition that inhibits *llgA* (7). *LlgA* is the late lytic gene activator that activates late phage genes. At 37°C and during *L. monocytogenes* intracellular infection, *llgA* is inhibited. Therefore, enhanced inhibition of *llgA* is a likely mechanism for phage production defects in ΔPDE. An established function of c-di-AMP is to regulate potassium (K^+^) homeostasis by inhibiting potassium import and activating potassium export. In addition, potassium has been shown to enhance lytic phage resistance in *Bacillus subtilis* (27). In *L. monocytogenes*, we found that potassium availability in the media did not impact spontaneous prophage induction, nor does it explain ΔPDE defect.

Approximately 8300 Listeria phages have been isolated and described to date (27). Lysogenic phages are a driver of bacterial physiology and population dynamics within a community. Prophage-encoded genes can determine bacterial host physiology and pathogenesis. Cholera toxin, Enterohemorrhagic *E. coli* Shiga toxin, and botulinum toxin are well-studied examples of bacterial virulence factors encoded within prophages (7). In addition, prophages also carry auxiliary metabolic genes that, despite dispensable for host growth, directly modulate bacterial metabolism and cell structure (7). Prophages often integrate in the host genome at sites of important genes, and thus, regulate the expression of these genes by excision and reintegration into this site (7). Under laboratory conditions, prophages can be induced by several stress conditions. DNA-damaging treatments, H_2_O_2_, pH fluctuations and fluoroquinolone and beta-lactam antibiotics are example stressors that activate prophage induction via the SOS response (28, 29). In addition, there are increasing examples of other inducers that act independently of the SOS response, such as various quorum sensing molecules and natural products (28, 29). Identification these signals and defining their activation mechanisms is essential to understand and predict how prophages can impact bacterial community dynamics.

## MATERIALS AND METHODS

### Strains and culture conditions

*L. monocytogenes* strains are listed in Table S1. Improved *Listeria* synthetic medium (LSM) was prepared based on the published recipe (7). Over-expression of genes was accomplished using the integrative plasmid pPL2 (7). Constructs were cloned in *XL1-Blue E. coli* with 25 μg/mL chloramphenicol and integrated into *L. monocytogenes* through conjugation with *SM10 E. coli*. Gene allelic exchange was performed using plasmid pLIM1.For all experiments related to overnight broth cultures, *L. monocytogenes* was grown at 37°C, with agitation, and for daytime cultures, at 30°C without shaking and with antibiotics, as needed. For glycerol stocks and maintenance, strains carrying derivatives of the integrative plasmid pPL2 were grown in brain heart infusion (BHI) broth with 200 µg/mL streptomycin and 10 µg/mL chloramphenicol.

### Bacterial lysis under Mitomycin-C treatment

Bacteria were grown overnight at 37°C, with agitation, in LSM broth and BHI broth. The overnight bacterial cultures were used to grow daytime cultures in LSM or BHI at 30°C, without shaking, to an OD at 600 nm (OD_600_) of 0.4, and then diluted to an OD_600_ of 0.15, which was then pipetted into a 96-well plate with or without Mitomycin-C (3 μg/mL). The plates were incubated at 30°C, without shaking, for 16h, in a Varioskan LUX Multimode Microplate Reader, and the OD_600_ was measured every 30 min. All experiments were repeated at least three times.

### Mitomycin-C treatment for phage quantification

Bacteria were grown overnight at 37°C, with agitation, in LSM or BHI broth, then diluted by a factor of 10 in fresh LSM or BHI broth, incubated without agitation at 30°C, to reach an OD_600_ of 0.4, then diluted to an OD_600_ of 0.15, and the lytic cycle was induced by the addition of Mitomycin-C (3 μg/mL) and incubation at 30°C for 16 h, without agitation. Bacterial cultures were passed through 0.22 μm filters that do not allow the passage of bacteria. These filtered supernatants were used to quantify phages through plaque assay

### Plaque assay

Dilutions of the filtered supernatants (mitomycin-C treated or from culture supernatants) were added to 3 mL melted LB-0.7% agar medium at 56°C, supplemented with 10 mM CaCl_2_, 10mM MgSO_4_, and 100 μL of 5×10^9^ CFU culture of *L. monocytogenes* MACK861 (used as indicator strains), and quickly overlaid on pre-warmed BHI-agar plates. Plates were incubated for 1 day in 30°C to allow plaques to form. All experiments were repeated at least three times.

### Co-culture assay

Bacterial cultures were grown overnight in LSM broth, at 37°C, with agitation. Overnight cultures were washed with 1x PBS and resuspended to OD_600_ 1, and then were used to inoculate fresh LSM media, at a dilution factor of 100, in equal ratios. The daytime cultures were incubated at 30°C, without agitation.

### Growth inhibition by phage addition

Bacterial cultures were grown overnight in LSM broth, at 37°C, with agitation. Overnight cultures were washed with 1x PBS and resuspended to OD_600_ 1, and then were used to inoculate fresh LSM media, at a dilution factor of 100, in equal ratios. The daytime cultures were incubated at 30°C, without agitation, and grown to OD_600_ 0.5. Phage φ10403S of MOI 0.1 was added to the cultures, and then the cultures were incubated at 30°C without agitation. Growth and phage production were quantified by OD_600_ measurement, CFU counting, and plaque assay at different time points.

### Quantification of reactive oxygen species

Bacterial cultures were grown in LSM for 12h, 16h and 24h, washed, and diluted to 5 × 10^8^ CFU/mL in 1x PBS. 200μL of diluted cultures were transferred into a black 96-well plate with 40 ng/μL of H_2_DCFDA (ThermoFisher). Fluorescence was measured in the Biotek Synergy H1 plate reader by excitation at 490 nm and emission at 530 nm over 30 minutes. Reactive oxygen species were quantified by the rates of increase in fluorescence intensity over 30 minutes.

