## Supplementary material for "*Listeria monocytogenes* prophage induction is activated by ppGpp and inhibited by c-di-AMP": Figures S1-S7

**Table S1:** Strains used in this study

| Strain number | Genotype | Source |
|---|---|---|
| ***L. monocytogenes*** (derivatives of strain 10403S) | | |
| TNHL261 | WT | (1) |
| TNHL325 | ΦWT | This study |
| TNHL22 | Δ*pdeA* Δ*pgpH* (ΔPDE) | (2) |
| TNHL493 | ΦΔPDE | This study |
| TNHL1257 | WT $P_{spac}$-*cI* | This study |
| TNHL1217 | ΦWT $P_{spac}$-*cI* | This study |
| TNHL1258 | ΔPDE $P_{spac}$-*cI* | This study |
| TNHL1218 | ΦΔPDE $P_{spac}$-*cI* | This study |
| TNHL1269 | WT $P_{spac}$-*kat* | This study |
| TNHL1260 | ΦWT $P_{spac}$-*kat* | This study |
| TNHL1259 | ΔPDE $P_{spac}$-*kat* | This study |
| TNHL1261 | ΦΔPDE $P_{spac}$-*kat* | This study |
| TNHL227 | Δ*relAPQ* | This study |
| TNHL680 | ΔPDE Δ*relAPQ* | This study |
| TNHL736 | Δ*relAPQ codY*::spec | This study |
| TNHL763 | ΔPDE Δ*relAPQ codY*::spec | This study |
| TNHL266 | WT l*acZ*::erm | This study |
| TNHL1119 | ΦWT *lacZ*::erm | This study |
| TNHL1219 | ΔPDE *lacZ::*erm | This study |
| TNHL1221 | ΦΔPDE *lacZ*::erm | This study |
| **Other *Listeria* strains** | | |
| TNHL269 | *Listeria monocytogenes* MACK861 | (3) |
| TNHL315 | *Listeria monocytogenes* F2365 | (4) |
| TNHL728 | *Listeria innocua* CLIP 11262 | (5) |
| ***E. coli*** | | |
| TNH1338 | pPL2-$P_{spac}$-*cI* in XL1B | This study |
| TNH1355 | pPL1-$P_{spac}$-*kat* in XL1B | This study |

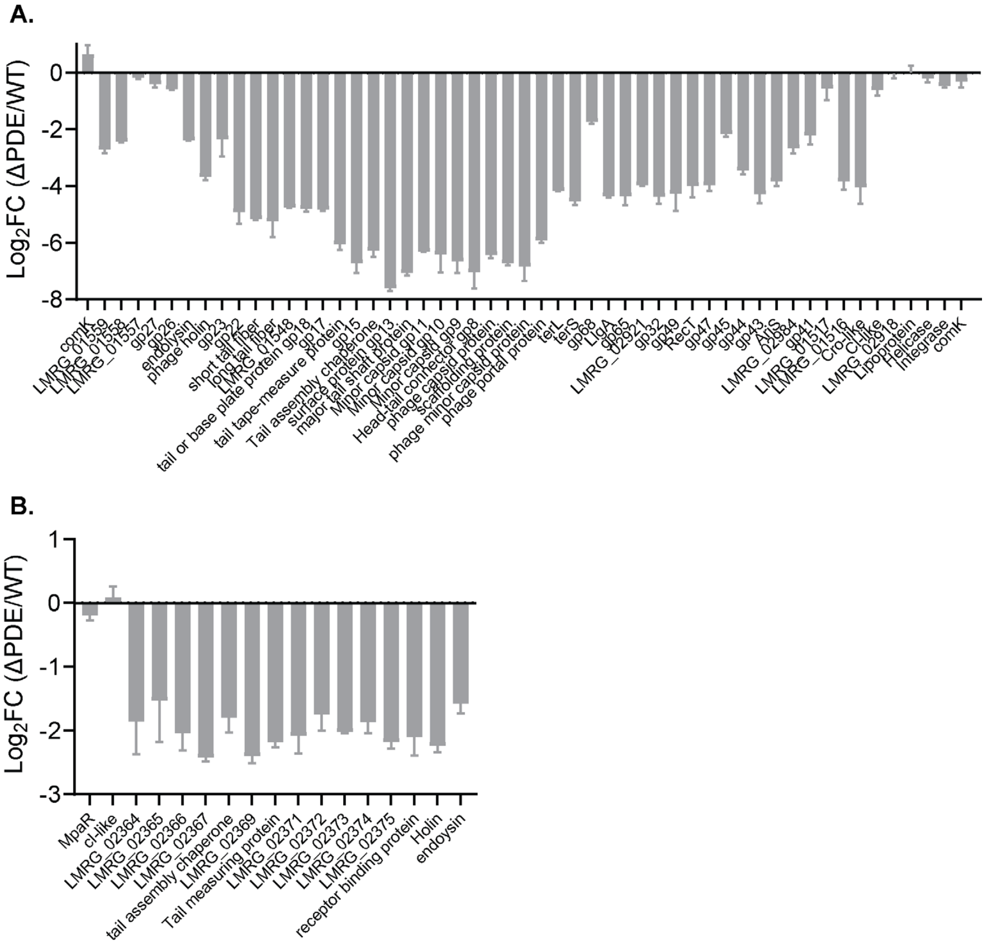

**Figure S1: Differentially expressed genes within the prophage and monocin loci of the ΔPDE strain.** RNAseq was performed on mid-log cultures grown in Listeria synthetic medium in 37°C. $Log_2$fold change of prophage and monocin gene expression was calculated from read counts per million of two independent ΔPDE cultures compared to the average of two independent WT cultures. Data in A and B are from the same RNAseq experiment. Error bars represent standard deviations.

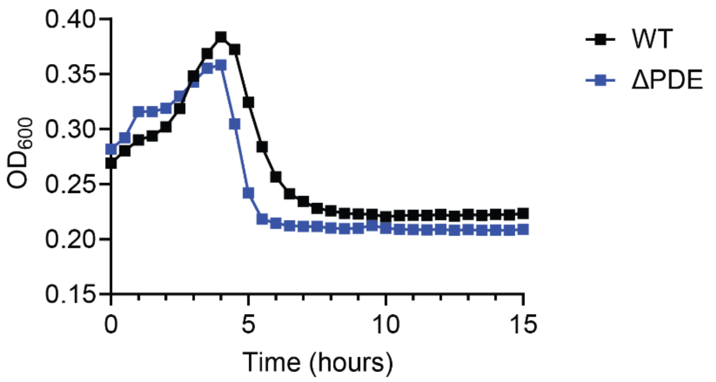

**Figure S2: Lysis of *L. monocytogenes* cultures following mitomycin-C treatment.** *L. monocytogenes* cultures were grown in LSM at 30°C to early log phase and treated with 3 μg/mL mitomycin-C. Culture density was monitored from the point of mitomycin-C addition.

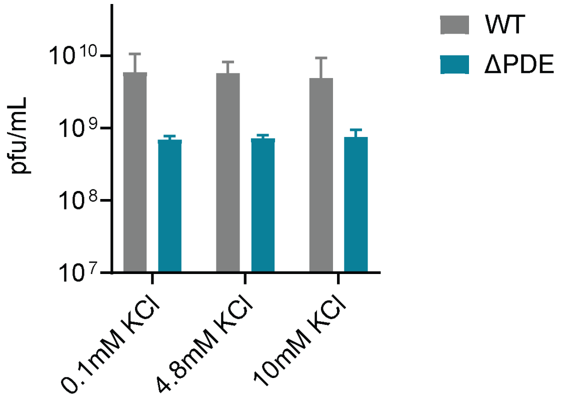

**Figure S3: Potassium does not impact prophage production by WT and ΔPDE.** *L. monocytogenes* was grown in LSM at 30°C to early log phase, then shifted to modified LSM with varying KCl and 3 μg/mL mitomycin-C. Culture supernatants were harvested at 16 hours after mitomycin-C treatment and quantified for Φ10403S phage particles using *Listeria monocytogenes* MACK861 as an indicator strain. Data are average of at least three independent experiments. Error bars represent standard deviations.

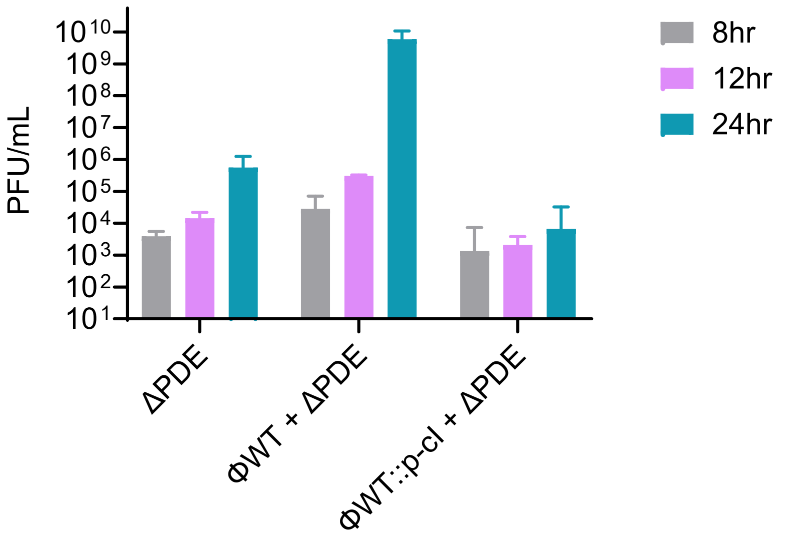

**Figure S4: Φ10403S phage production is amplified in late co-cultures.** *L. monocytogenes* was grown in LSM in co-cultures with 1:1 inoculation at 30°C and at indicated time points, culture supernatants were collected for quantifying Φ10403S phage production using *Listeria monocytogenes* MACK861 as an indicator strain. Data are average of at least three independent experiments. Error bars represent standard deviations.

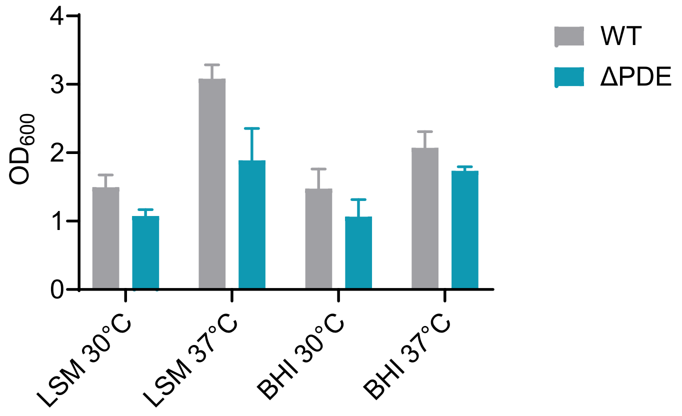

**Figure S5: *L. monocytogenes* growth.** *L. monocytogenes* was grown in BHI and LSM at 30°C statically and 37°C shaking for 24 hours, reaching the indicated $OD_{600}$, at which culture supernatants were harvested to quantify Φ10403S phage particles. Data are average of at least three independent experiments. Error bars represent standard deviations.

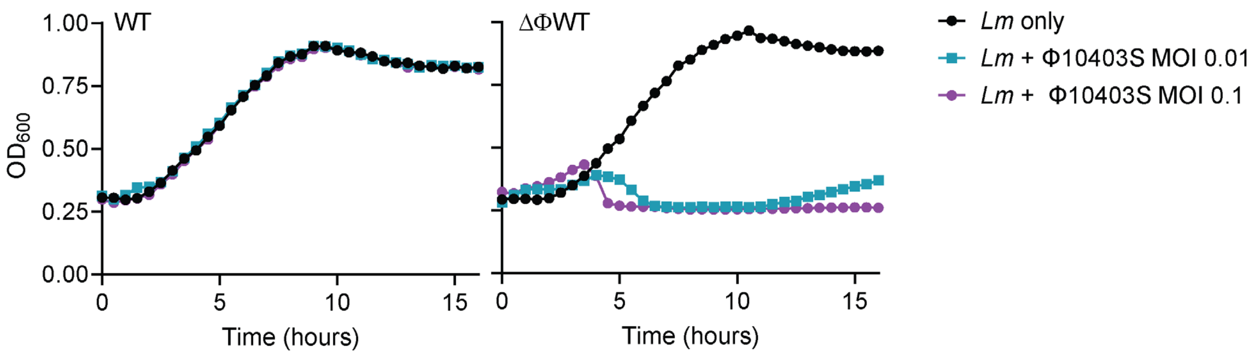

**Figure S6: Φ10403S lysogen protects *L. monocytogenes* from super-infection.** Phage Φ10403S, harvested from ΔΦWT + ΔPDE co-cultures, were used to infect *L. monocytogenes*, grown in LSM at 37°C, at indicated multiplicity of infection (MOI). Bacterial growth post infection was monitored by $OD_{600}$.

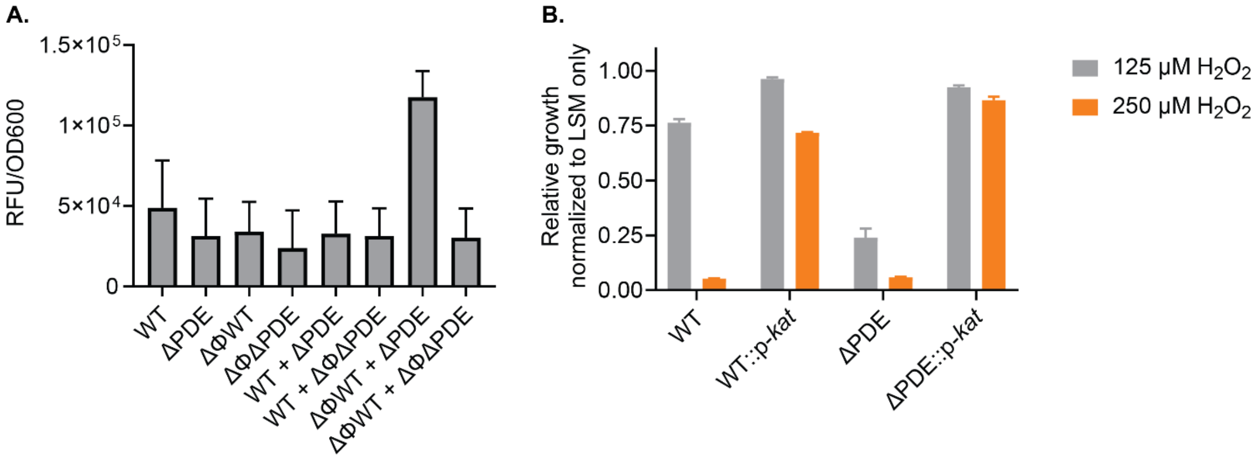

**Figure S7: $H_2O_2$ contributes to Φ10403S induction. A.** Reactive oxygen species quantified using $H_2$DFCA, in *L. monocytogenes* cultures grown in LSM at 30°C for 24 hours. **B.** Over-expression of *kat* increases $H_2O_2$ resistance. *L. monocytogenes* cultures were grown with shaking in LSM with varying $H_2O_2$ concentrations for 16 hours. For each strain, $OD_{600}$ of cultures grown in $H_2O_2$ were normalized to $OD_{600}$ of the same strain grown in LSM only. Data are average of at least three independent experiments. Error bars represent standard deviations.
